# Interference of feral radish (*Raphanus sativus*) resistant to AHAS-inhibiting herbicides in oilseed rape, wheat and sunflower crops

**DOI:** 10.1101/2020.08.27.270942

**Authors:** Roman B. Vercellino, Claudio E. Pandolfo, Miguel Cantamutto, Alejandro Presotto

## Abstract

*Raphanus sativus* (feral radish), a cosmopolitan weed, has developed resistance to acetohydroxyacid synthase (AHAS) inhibitor herbicides in several countries of South America. This study reports the effects of season-long interference of several feral radish densities on grain yield and yield components of oilseed rape, wheat and sunflower, and on feral radish traits under field conditions. Feral radish density treatments consisted of 0, 2, 4, 8 and 16 plants m^−2^ in oilseed rape, 0, 4 and 12 plants m^−2^ in wheat, and 0, 1.6, 4, 8 and 16 plants m^−2^ in sunflower. The number of inflorescences per area, seeds per inflorescence and the seed biomass of crops were reduced with increasing feral radish densities. The rectangular hyperbola model revealed yield losses by up to 100 %, 74.4 % and 12.2 % in oilseed rape, wheat and sunflower, respectively. Feral radish seed production ranged from 4,300 to 31,200, and 1000 to 4,700 seeds m^−2^ in winter crops and sunflower, respectively. Season-long feral radish interference can result in serious economic losses in oilseed rape, wheat and sunflower. The adverse impact of feral radish on the yield of winter and summer crops and the high feral radish seed and pods production suggests the need for the development and implementation of diverse and effective long-term weed management practices.

## Introduction

Weeds are considered the major constraint to food production in agricultural systems around the world (Oerke 2006). Chemical weed management is currently the most adopted and the most effective approach for controlling weeds all over the world (Bajwa 2014, Scursoni et al. 2019). Repeated applications of herbicides in arable fields have resulted in the rapid evolution of resistant populations in at least 262 weed species worldwide (Heap 2020), possibly favoured by the increasing adoption of the no-till system together with a reduction in crop rotations (Kraehmer et al. 2014). Understanding the potential crop loss from herbicide resistant weeds provides useful information for developing and/or optimizing integrated weed management strategies and understanding the strengths and weaknesses of target weeds.

*Raphanus sativus* L. (radish) is an ancient root crop from the Brassicaceae family. Weedy radish populations are considered as de-domesticated (feral) forms derived from the cultivated radish. Feral radish grows naturally in Europe, North America, South America and Japan, and it is a problematic winter weed in temperate zones of the Americas (Snow and Campbell 2005, Theisen 2008, Vercellino et al. 2018). Its probable wild ancestor, *Raphanus raphanistrum* (wild radish), is a successful invader and is one of the worst weeds worldwide (Snow and Campbell 2005). In Australia, the southern United States and southern Brazil, *R. raphanistrum* is considered one of the most troublesome weeds in winter crops (Lamego et al. 2013, Snow and Campbell 2005, Warwick and Francis 2005). Both species have developed resistance to herbicides, including resistance to acetohydroxyacid synthase (AHAS) inhibitor herbicides (Heap 2020). AHAS-resistant feral radish has been found in Brazil, Chile and Argentina (Heap 2020, Pandolfo et al. 2016, Theisen 2008), and wild radish in Australia, Brazil and South Africa (Heap 2020). Feral radish and wild radish are self-incompatible and insect-pollinated with an annual or biennial life cycle (Snow and Campbell 2005). High seed production, a darkness requirement for germination, the presence of a pericarp covering the seeds, and seed germination in a broad range of temperatures are some of attributes that favour the emergence and establishment of feral radish (Vercellino et al. 2018, 2019), as in the wild relative *R. raphanistrum* (Cheam 1986, Malik et al. 2010).

Feral radish is well distributed throughout the Argentinean agricultural landscape and is one of the most noxious weeds in the southeast of Buenos Aires province, the main winter cereal producing area (Scursoni et al. 2014). In that area, we reported the occurrence of several AHAS-resistant feral radish populations infesting winter and summer crops (Pandolfo et al. 2013, 2016, Vercellino et al. 2018). Since the initial report in 2011, the spread of feral radish resistant to AHAS herbicides has rapidly increased, and currently, its presence has been reported in Buenos Aires, Tucumán and Salta provinces (AAPRESID 2019).

AHAS-resistant feral radish must be controlled in both winter and summer crops to prevent yield losses. In Argentina, since more than 90 % of agricultural area is sown under no-till system (ReTAA 2020), weed control depends primarily on herbicides (Scursoni et al. 2019). In some crops, pre-emergence herbicides may provide effective control of AHAS-resistant feral radish (Pandolfo et al. 2016). On the other hand, post-emergent broadleaf weed control in broadleaf crops is not easy. Farmers have a few (e.g. aclonifen) or no herbicide options for post-emergence control of AHAS– resistant feral radish in sunflower and oilseed rape crops, respectively (Pandolfo et al. 2016). In winter cereals (i.e. wheat and barley), post-emergent control of AHAS-resistant feral radish is limited to phenoxy (2,4-D and MCPA) and photosystem-II inhibitor (bromoxynil) herbicides. For these reasons, several extensive populations of AHAS-resistant feral radish, due to the Trp574Leu mutation, have been found severely infesting both winter and summer crops (Pandolfo et al. 2016, Theisen 2008, Vercellino et al. 2018).

Limited information exists on the effect of feral radish interference on both winter and summer crops. In one study in Brazil, feral radish interference at densities of >40 plants m^−2^ reduced soybean yield by 15 % (Bianchi et al. 2011). Interference studies on AHAS-resistant feral radish in oilseed rape, wheat and sunflower will provide useful information on the importance of controlling this species using integrated weed management strategies. The aims of this study were: (1) to estimate the impact of the interference of several densities of AHAS-resistant feral radish on the yield and yield components of oilseed rape, wheat and sunflower crops, and (2) to estimate feral radish dry biomass and seed production at several densities in oilseed rape, wheat and sunflower crops. Feral radish densities were 0, 2, 4, 8 and 16 plants m^−2^ in oilseed rape, 0, 4 and 12 plants m^−2^ in wheat, and 0, 1.6, 4, 8 and 16 plants m^−2^ in sunflower. This is the first report of feral radish interference in oilseed rape, wheat and sunflower.

## Material and methods

### Plant material

Two AHAS-resistant feral radish populations (RSBA10 and RSBA3-R) carrying the Trp574Leu mutation, originating from southern Buenos Aires province (Argentina) and previously characterized by Pandolfo et al. (2016) and Vercellino et al. (2018), were used in these studies. RSBA10, with 95 % of individuals resistant to AHAS herbicides carrying the Trp574Leu mutation, was collected in an IMI-resistant sunflower field treated with imazapyr near Pieres, Lobería county (S 38° 24’, W 58° 35’). RSBA3 was collected in an IMI-resistant oilseed rape field treated with imazethapyr (RSBA3) near Balcarce (S 37° 35’, W 58° 31’). A purified homozygous population with the AHAS Trp574Leu mutation (RSBA3-R) was generated from RSBA3 by selecting 15 homozygous resistant plants using a specific Trp574Leu mutation CAPS marker and following the procedure described in Pandolfo et al. (2016). In a previous study, no differences were found in the phenology and pods and seeds production between RSBA10 and RSBA3 (Vercellino et al. 2018). We generally used the RSBA10 accession in interference experiments.

### Field experiments

Field experiments evaluating feral radish interference in oilseed rape, wheat and sunflower were conducted between 2016 and 2018 in the Agronomy Department experimental field at the Universidad Nacional del Sur, Bahía Blanca, Argentina (38°41’46’’S, 62°14’55’’W). The site is characterized as semi-arid with a temperate climate, with a mean annual temperature of 14.9 °C; the mean temperatures of the coldest (July) and the hottest (January) month are 7.6 and 23.0 °C, respectively. Average annual precipitation is 641 mm (National Meteorological Service 2019, https://www.smn.gob.ar/). The soil has a well-drained loamy sand texture with 1.1 % organic matter and pH 7.7 (a typical soil in the southwest of Buenos Aires province). Experiments were drip irrigated as necessary for optimal growing conditions.

Under farm conditions, we observed the emergence of feral radish in the planting row probably because the seeder disc broke the pods and released and buried their seeds during cultivation, as reported by Cheam (1986) in the wild relative *R. raphanistrum*. For this reason, we evaluated the interference of feral radish established at the same time as the crops. We selected the weed densities according to our previous surveys under field conditions. In each experiment, the target feral radish densities were achieved by manually sowing 2–3 seeds per hole and 2–3 cm deep, spaced according to the density plan. At the four–leaf stage of each crop (i.e. oilseed rape, wheat and sunflower), the target weed stand was adjusted by hand thinning. Hand weeding was carried out periodically to avoid interference from other weeds. Three different set of experiments were performed:

#### a. Oilseed rape

The oilseed rape cultivar Hyola 575 CL was planted by hand in 0.2-m rows at 50 plants m^−2^ on May 2017 and 2018, during the winter growing season. Experimental units (plots) were seven rows wide spaced at 0.2 m by 2 m long. Simultaneously, feral radish seeds were planted, as was described above, to achieve 0 (control), 2, 4, 8 and 16 plants m^−2^. Experiments were fertilized with 90 kg ha^−1^ diammonium phosphate at planting and 200 kg ha^−1^ urea at the four–six-leaf stage. The experimental design was a randomized complete block with four replications. The experiments were performed in the same manner in 2017 and 2018.

At oilseed rape maturity, the plant height, number of branches, and pods per main inflorescence were measured in ten successive oilseed rape plants in the middle of the three central rows of each experimental unit. Then, the plants were hand-harvested and dried at 60 °C to constant weight to homogenize treatments. After that, we quantified seeds per pod, seed weight and yield per plant. Seeds per pod was estimated by averaging the number of seeds per pod of five pods per plant. Each plant was threshed by hand and its seeds were cleaned and weighed to obtain the plant yield. The seed weight was estimated by averaging the weight of three replicates of 100 seeds per plant. Data from the ten oilseed rape plants per experimental unit were averaged for the statistical analysis. Oilseed rape seed weight and yield were standardized to 8 % moisture content.

#### b. Wheat

Wheat cultivars ACA 360 and KLEIN PROTEO were planted in mid-May 2018 (winter) and in early August 2018 (spring), respectively. Crop density was adjusted to 200 and 350 plants m^−2^ for winter and spring wheat, respectively. Plots were nine rows wide spaced at 0.2 m by 1.5 m long. Simultaneously, feral radish seeds were planted, as was described above, to achieve 0 (control), 4 and 12 plants m^−2^. Experiments were fertilized with 90 kg ha^−1^ diammonium phosphate at planting and 200 kg ha^−1^ urea at the five–leaf stage. The experimental design was a randomized complete block with four replications. The experiments were performed in the same way in both growing seasons.

At wheat maturity, plant height was measured on three replicates of five successive plants, and three replicates of 0.5 m long in the middle of the three central rows of each experimental unit were hand-harvested and dried at 60 °C to constant weight to homogenize treatments and the growing seasons. In each replicate, we also evaluated the number of spikes m^−2^, spikelets per spike, grains per spikelet, grain weight and yield. Spikelets per spike and grains per spikelet were obtained by averaging five spikes, and grain weight by averaging the weight of four replicates of 100 grains.

Data from wheat plants of each experimental unit were averaged for the statistical analysis. Wheat grain weight and yield were standardized to 13.5 % moisture content.

#### c. Sunflower

The sunflower cultivar SYN 3970 CL was planted at 6.4 plants m^−2^ on Nov 2016, during the summer growing season. Plots were five rows wide spaced at 0.52 m by 2.1 m long. Simultaneously, feral radish seeds were planted, as was described above, to achieve 0 (control), 1.6, 4, 8 and 16 plants m^−2^. The experimental plots were fertilized with 90 kg ha^−1^ diammonium phosphate at planting and 150 kg ha^−1^ urea at the four–six-leaf stage. Experimental design was a randomized complete block with four replications.

At the end of crop flowering, the plant height, width of the three middle leaves and the number of green leaves were measured on three successive sunflower plants in the middle of the central row of each experimental unit. The leaf area was estimated according to Aguirrezábal et al. (1996) from the leaf width, and the leaf area was obtained by averaging the leaf area of three leaves per plant. At sunflower maturity, the heads of the three evaluated plants were hand-harvested and dried at 60 °C to constant weight. After that, we measured the head diameter, seeds per plant, seed weight and yield per plant in the three plants harvested per experimental unit. The seed weight was estimated by averaging the weight of four replicates of 100 seeds. Data from the sunflower plants in each experimental unit were averaged for the statistical analysis. Sunflower seed weight and yield were standardized to 11 % moisture content.

### Weed traits

At the experimental level, we measured the plant height in four successive feral radish plants located in the centre of each experimental unit at weed maturity. Then, plants were hand-harvested, dried at 60 °C to constant weight and weighed to determine the dry biomass per weed plant. After that, pods from these plants were counted and crushed by hand or using a mortar. Seeds were cleaned and weighed to obtain the plant yield. The seed weight was estimated by averaging the weight of three replicates of 100 seeds per plant. The number of seeds per plant was estimated by dividing the plant yield by the seed weight for each plant. Seeds per pod was estimated by dividing the seeds per plant by the number of pods per plant. Data from the feral radish plants per experimental unit were averaged for the statistical analysis. Weed traits were measured in the same way for all three different set of experiments.

### Statistical analysis

The relationship between the crop yield and feral radish density was analysed for each crop and year (i.e. oilseed rape) or growing season (i.e. wheat) with the rectangular hyperbolic model (Cousens 1985) using PROC NLIN procedure in SAS/STAT software (SAS University Edition, SAS Institute Inc., Cary, NC, USA).

The yield loss relationship is described by Equation 1:

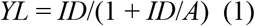

where *YL* is the percentage crop yield loss relative to the weed-free treatment, *I* is the percentage crop yield loss per weed as the weed density approaches zero, *D* is the weed density, and *A* is the percentage crop yield loss as the weed density approaches infinity. The model was constrained to an asymptotic percentage yield loss, A lying between 0 and 100% because yield loss cannot exceed 100%. The parameter *I* is used as an indicator of potential weed competitiveness (Cousens 1985).

As no other crop data fitted a logical/appropriate regressions model, crop data were analysed using general linear models (GLMs) with PROC GLM in SAS. Feral radish density and year (i.e. oilseed rape) or growing season (i.e. wheat) were considered as fixed factors, to test for significant main effects and interactions. ANOVA and Fisher’s Protected LSD tests were used to determine the variation between treatments.

To analyse the relationships between feral radish traits (e.g. dry biomass, pod number and seed production) and plant density for each experiment, we performed regression analyses using Prism software (Graphpad Prism 7.0; GraphPad Software, San Diego, California, USA). These relationships were established from the best fits of the experimental data to the appropriate functions, and the coefficients of determination (R^2^) were reported. ANOVA was used to determine the variation between treatments in weed plant height, seeds per pod and seed weight.

The two-parameter hyperbolic regression model (Equation 2) was used to describe the density-dependent effects of feral radish on the dry biomass of feral radish, the pod number and seed production per meter square:

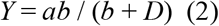

where *Y* is feral radish dry biomass, pod number and seed production per square meter, *a* is the asymptote or estimate of maximum feral radish dry biomass, pod number and seed production, *b* is the estimate of the feral radish density at which 50 % of maximum feral radish dry biomass, pod number and seed production, and *D* is weed density.

The one-phase decay regression model (Equation 3) was used to describe the density-dependent effects of feral radish on feral radish dry biomass, pod number and seed production per plant:

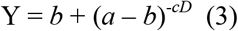

where *Y* is feral radish dry biomass, pod number and seed production per plant, *b* is the Y value at infinite feral radish density, expressed in the same units as Y, *a* is the Y value when D approaches zero, and *D* is weed density.

## Results and Discussion

### Oilseed rape

In both years, the oilseed rape yield decreased substantially as the feral radish density increased. Analysis of oilseed rape traits revealed no significant interactions between weed density and year in the plant height, number of branches, pods per main inflorescence and seeds per pod, therefore data from the two growing seasons were pooled (Table 1 and S1). However, there was a significant interaction in the seed weight, therefore this data was analysed separately by year (Table 1 and S1). Feral radish interference had a significant effect on plant height, number of branches, pods per main inflorescence and seeds per pod (Table 1 and S1). The plant height was reduced at 4 feral radish m^−2^ and increasing weed density from 4 to 16 plants m^−2^ reduced the plant height by up to 15.6 %. The number of branches was reduced at 2 weeds m^−2^ and increasing feral radish density from 2 to 16 plants m^−2^ decreased the number of branches by up to 35.7 % (Table 1). Pods per main inflorescence and seeds per pod were only reduced at the high weed density (16 plants m^−2^), and the reduction was 20.6 and 18.4 %, respectively (Table 1). Similarly, the number of branches, the pods per inflorescence and the seeds per pod were the most affected yield components in oilseed rape during interference in several studies (Rondanini et al. 2017, Holman et al. 2004). Seed weight was not affected by feral radish interference in 2017, but there were significant reductions (19.8 to 28.5%) at the densities ≥ 8 feral radish m^−2^ in 2018 (Table 1).

**Table 1.** Oilseed rape traits under interference of five feral radish densities

| Weed<br>$\text{m}^{-2}$ | Plant height<br>(cm) | Number of<br>branches | Pods per main<br>inflorescence | Seeds per pod | Seed biomass<br>(mg) 2017 | Seed biomass<br>(mg) 2018 |
| --- | --- | --- | --- | --- | --- | --- |
| 0 | 99.2 ± 4.4 a | 5.86 ± 0.24 a | 27.6 ± 1.4 ab | 17.4 ± 0.4 ab | 3.27 ± 0.06 | 3.46 ± 0.08 a |
| 2 | 97.6 ± 5.9 ab | 5.30 ± 0.18 b | 29.2 ± 2.0 a | 17.8 ± 0.4 a | 3.27 ± 0.16 | 3.25 ± 0.13 a |
| 4 | 91.8 ± 3.9 bc | 4.56 ± 0.30 c | 26.6 ± 1.0 ab | 17.4 ± 0.3 ab | 3.20 ± 0.05 | 3.22 ± 0.11 a |
| 8 | 92.4 ± 5.0 ab | 4.66 ± 0.21 c | 25.1 ± 1.6 b | 16.1 ± 0.4 b | 3.05 ± 0.12 | 2.77 ± 0.14 b |
| 16 | 84.9 ± 3.2 c | 3.77 ± 0.21 d | 21.9 ± 1.4 c | 14.2 ± 0.5 c | 3.12 ± 0.10 | 2.46 ± 0.02 c |
| ANOVA | ** | ** | ** | ** | ns | ** |

Observations are means ± SE of two years, except for the variable “seed weight” with showed density x year interaction, therefore means ± SE are shown for each year (2017 and 2018). For each trait, a different letter indicates significant differences (\* $P \leq 0.05$ Fisher’s LSD; \*\* $P \leq 0.01$ Fisher’s LSD).

The adjusted rectangular hyperbolic functions (R^2^ = 0.95 and 0.93) showed that feral radish decreased the oilseed rape yield per unit by 16.8 – 15.7 % at low weed density (parameter *I*), and by 100 % when the density approaches infinity (parameter *A*) for 2017 (*F* = 622.87; *P* < 0.0001) and 2018 (*F* = 374.18; *P* < 0.0001), respectively (Fig. 1). Oilseed rape yields with the weed-free treatment were 6706 ± 340 and 4647 ± 302 kg ha^−1^ in 2017 and 2018, respectively. Differences observed between years were probably related to prevailing environmental conditions during the growing season. Despite the differences in oilseed rape yield between the two years, oilseed rape yield losses were very similar (Fig. 1). Our results are similar to those obtained by Blackshaw et al. (2002) in the wild relative *R. raphanistrum*, although they used a wider range of weed densities (0 to >60 wild radish plants m^−2^), which could indicate that feral radish is more competitive than wild radish. High level of interference in oilseed rape was also found as a result of the interference of the Brassicaceae species *Sinapis arvensis* (Naderi and Ghadiri 2011). Because seeds of *R. sativus* have about 32.6 % erucic acid (Mandal et al. 2002) and 149 μM g^−1^ glucosinolates (Daxenbichler et al. 1991), seeds of feral radish could also reduce oilseed rape quality by increasing the erucic acid content of oilseed rape oil and the glucosinolate levels in oilseed rape meal that is marketed as livestock feed, as demonstrated in *R. raphanistrum* (Blackshaw et al. 2002).

**Figure 1.**
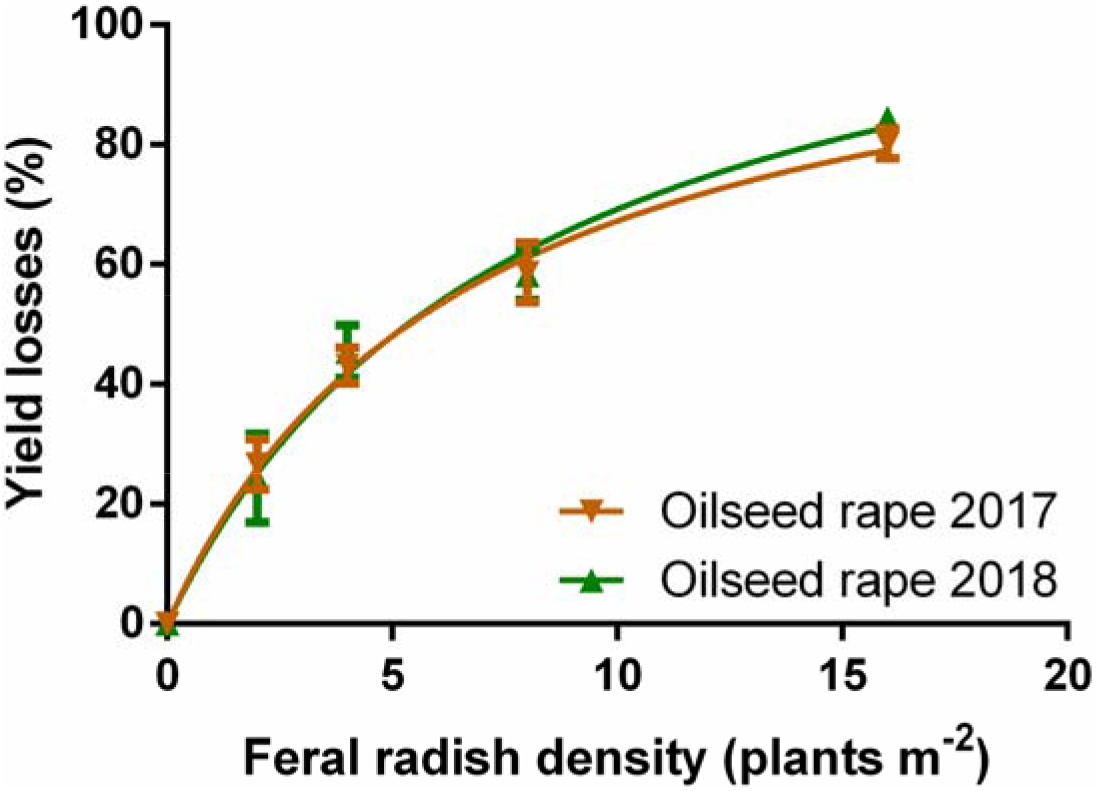
Relationship between feral radish density (plants m^−2^) and yield loss (%) of oilseed rape (2017 and 2018) crops described with a rectangular hyperbola model. Vertical bars indicate means ± SE. Data fit the model *YL* = 16.82D/(1 + 16.82D/112.10) (oilseed rape 2017) and *YL* = 15.72D/(1 + 15.72D/123.70) (oilseed rape 2018).

### Wheat

In both growing seasons, as the feral radish density increased the wheat yield also decreased considerably. Analysis of the wheat traits revealed no significant interaction between the weed density and growing season in the plant height, spikes m^−2^, spikelets per spike, grains per spikelet and grain weight, therefore data from the two growing seasons were pooled (Table 2 and S2). Interference from feral radish did not affect wheat plant height. However, spikes m^−2^, spikelets per spike, grains per spikelet and the grain weight were significantly reduced at 4 weeds m^−2^, and increasing feral radish density from 4 to 12 plants m^−2^ reduced the spikes m^−2^ by up to 36.3 %, the spikelets per spike by up to 7.5 %, the grains per spikelet by up to 20.1 % and the grain weight by up to 12.3 % (Table 2). Similar results were observed due to the interference of the grass weeds, *Avena fatua* and *Lolium persicum* (Holman et al. 2004, Guillen-Portal et al. 2006). However, the Brassicaceae species *Rapistrum rugosum* reduce wheat yield mainly due to a reduction in the number of spikes per area and grains per spike but, unlike our results, the weed did not affect the grain weight (Manalil and Chauhan 2019).

**Table 2.**
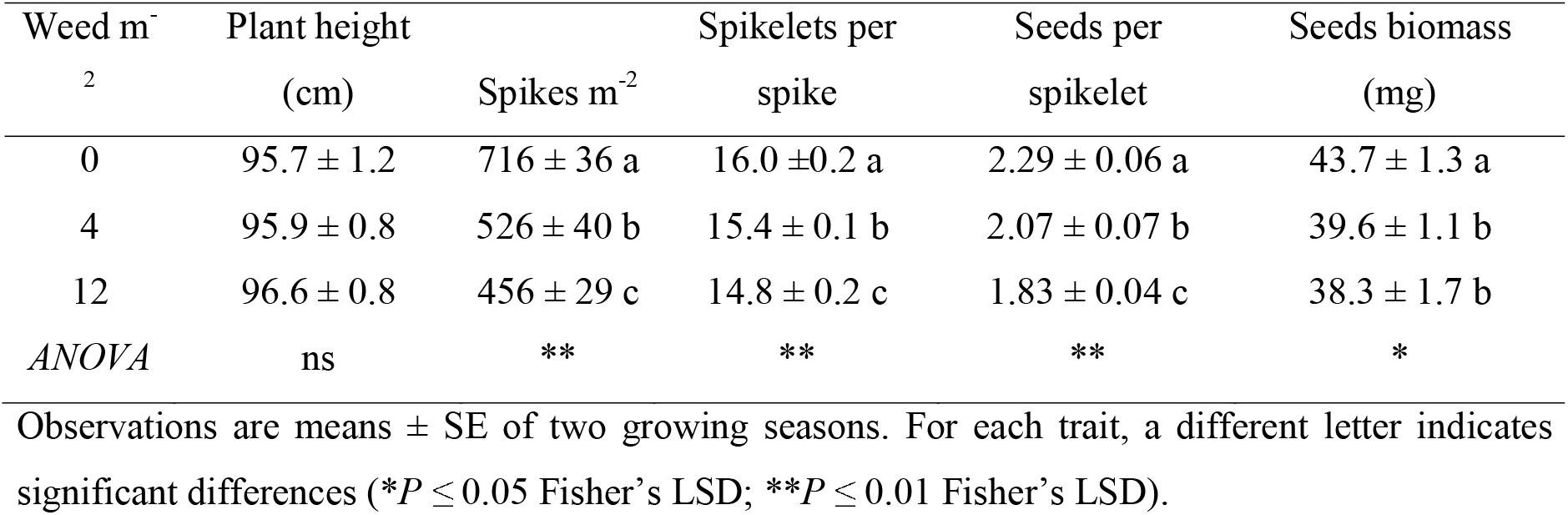
Wheat traits under interference of three feral radish densities.

| Weed m<sup>-2</sup> | Plant height<br>(cm) | Spikes m<sup>-2</sup> | Spikelets per<br>spike | Seeds per<br>spikelet | Seeds biomass<br>(mg) |
| --- | --- | --- | --- | --- | --- |
| 0 | 95.7 ± 1.2 | 716 ± 36 a | 16.0 ± 0.2 a | 2.29 ± 0.06 a | 43.7 ± 1.3 a |
| 4 | 95.9 ± 0.8 | 526 ± 40 b | 15.4 ± 0.1 b | 2.07 ± 0.07 b | 39.6 ± 1.1 b |
| 12 | 96.6 ± 0.8 | 456 ± 29 c | 14.8 ± 0.2 c | 1.83 ± 0.04 c | 38.3 ± 1.7 b |
| ANOVA | ns | ** | ** | ** | * |

Observations are means ± SE of two growing seasons. For each trait, a different letter indicates significant differences (\* $P \leq 0.05$ Fisher's LSD; \*\* $P \leq 0.01$ Fisher's LSD).

The adjusted rectangular hyperbolic functions (R^2^ = 0.93 and 0.86) showed that feral radish reduced the wheat yield per unit by 46.3 and 14.5 % at low weed density (parameter *I*), and by 62.2 and 74.4 % when the density approached infinity (parameter *A*) for winter (*F* = 204.15; *P* < 0.0001) and spring (*F* = 83.83; *P* < 0.0001) wheat, respectively (Fig. 2). Although, the low number of weed densities could result in over or underestimation of the parameter *I* (Fig. 2), the use of this model allowed us to have an estimation of the effect of feral radish interference on wheat yield at mid or high densities (situation commonly found under field conditions because weeds grow in patches) (Fig. 2). Wheat yields with the weed-free treatment were 8049 ± 299 and 8538 ± 204 kg ha^−1^ in winter and spring growing season, respectively. Differences observed between growing seasons were probably related to differences in the competitive capacity of wheat cultivars and prevailing environmental conditions during the growing season. Our results were similar to those obtained by Eslami et al. (2006) and Manalil and Chauhan (2019) in *R. raphanistrum* and *R. rugosum*, respectively, although they used a wider range of weed densities (0 to >60 wild radish plants m^−2^ and 0 to >45 *R. rogosum* plants m^−2^), which could indicate that feral radish is more competitive than wild radish and *R. rugosum*, two important Brassicaceae weeds.

**Figure 2.**
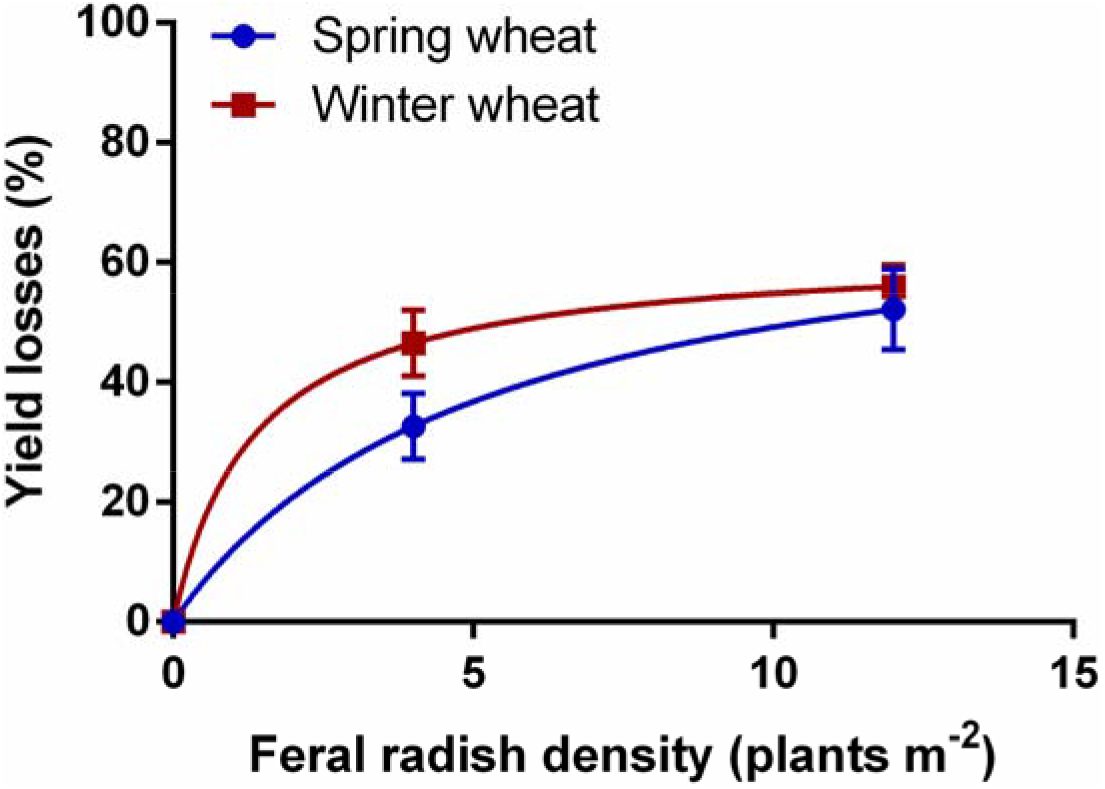
Relationship between feral radish density (plants m^−2^) and yield loss (%) of wheat (winter and spring) crops described with a rectangular hyperbola model. Vertical bars indicate means ± SE. Data fit the model *YL* = 46.26D/(1 + 46.26D/62.19) (winter wheat) and *YL* = 14.54D/(1 + 14.54D/74.44) (spring wheat).

### Sunflower

Sunflower yield was also affected by feral radish interference. The interference of feral radish reduced the sunflower plant height, green leaves per plant and head diameter. The plant height and number of green leaves were reduced by 6.5 and 11.0 %, only at the high weed density (16 plants m^−2^). The head diameter was reduced by 7.2% with 8 feral radishes m^−2^ (Table 3 and S3). However, the interference of feral radish did not significantly affect the number of seeds per head and the seed weight (Table 3 and S3). The rectangular hyperbolic functions (R^2^ = 0.57) showed that feral radish decreased (*F* = 48.36; *P* < 0.0001) the sunflower yield per unit by 5.4 % at low weed density (parameter *I*), and by 12.2 % when the density approached infinity (parameter *A*) (Fig. 3). The estimated weed-free sunflower yield was 3728 ± 236 kg ha^−1^. A similar result was found in the season-long interference of feral radish on soybean (Bianchi et al. 2011). The higher interference caused by feral radish in oilseed rape and wheat crops compared to sunflower and soybean (Bianchi et al. 2011) may possibly be because feral radish is a winter weed (Snow and Campbell 2005), and therefore it had larger plant size in winter than in summer (see below) (Vercellino et al. 2018). In another conditions, Pandolfo (2016) observed sunflower yield losses greater than 70% due to feral radish interference at farm level. This indicate that environmental conditions, as well as other factors (e.g. soil type and the timing of weed emergence relative to the crop), can play a key role in weed interference (Swanton et al. 2015).

**Table 3.**
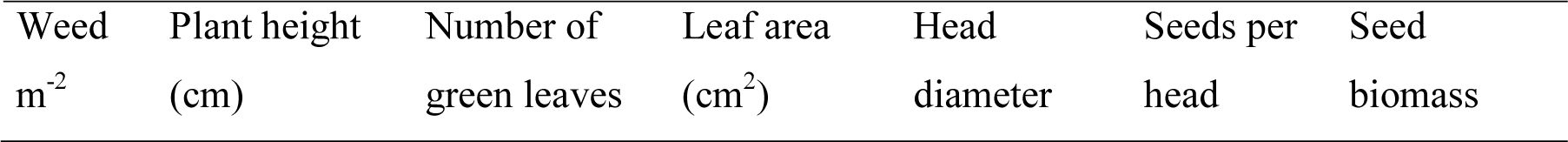
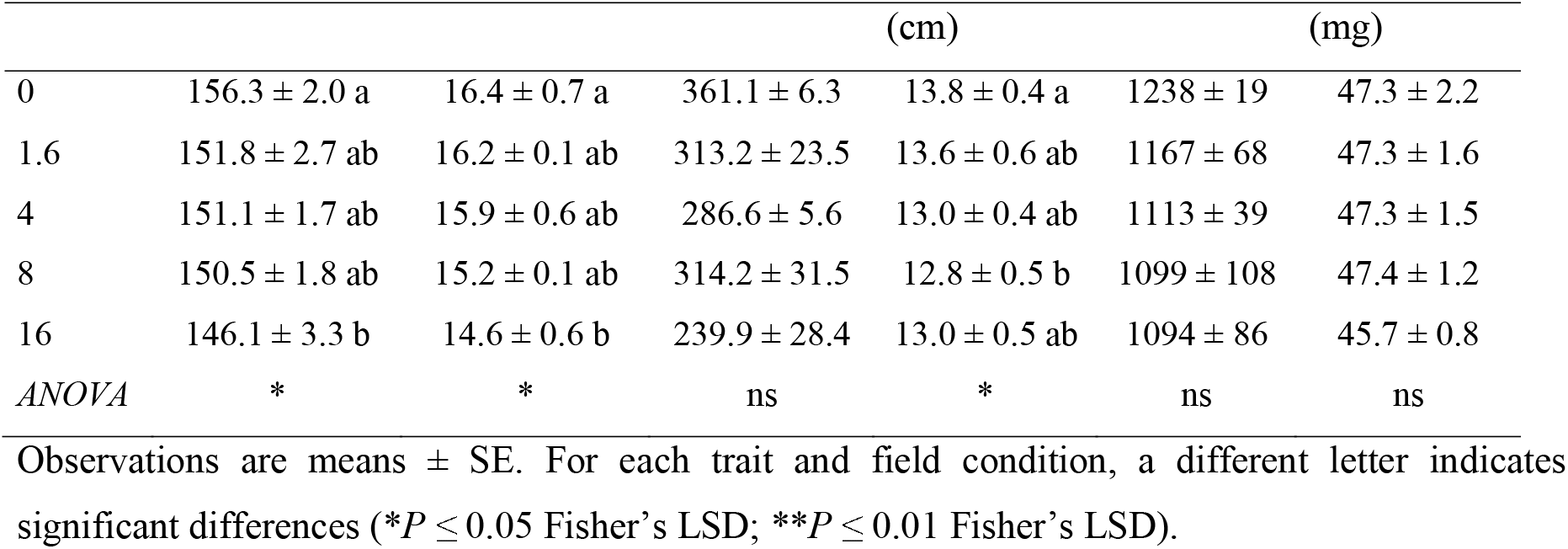
Sunflower traits under interference of feral radish under experimental and farm conditions.

**Figure 3.**
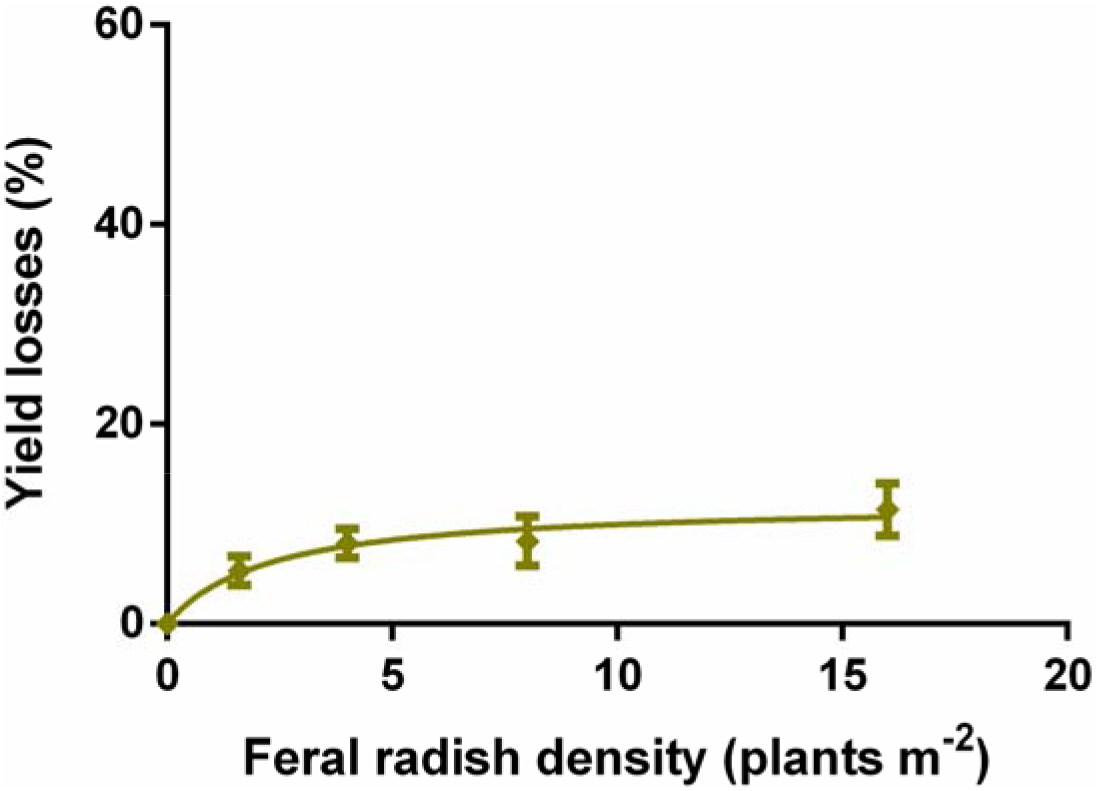
Relationship between feral radish density (plants m^−2^) and yield loss (%) of sunflower crop described with a rectangular hyperbola model. Vertical bars indicate means ± SE. Data fit the model *YL* = 5.43D/(1 + 5.43D/12.19).

Clearly, high-yielding oilseed rape, wheat and sunflower crops are sensitive to interference from a competitive broadleaf weed such as feral radish. These results showed that oilseed rape, wheat and sunflower yields tended to decrease as the feral radish density increased. In general, feral radish interference mainly reduced the number of inflorescences per area (represented by spikes m^−2^ in wheat and the number of primary branches by plant density in oilseed rape), followed by a reduction in the inflorescence size and the number of seeds per inflorescence (represented by spikelets per spike and by grains per spikelet in wheat, pods per inflorescence and by seeds per pod in oilseed rape, and head diameter and seeds per head in sunflower), and finally the seed weight. These results are in agreement with those previous that evaluate the effect of the weed interference in oilseed rape, wheat and sunflower crops (Manalil and Chauhan 2019, Guillen-Portal et al. 2006, Holman et al. 2004).

### Feral radish traits

All feral radish in these studies assumed an annual life cycle. Feral radish traits were more affected in the sunflower experiment than in either the oilseed rape or wheat experiments. The feral radish plant height was not affected by weed density, which ranged from 118.5 ± 7.9 cm to 149.0 ± 3.3 cm in both the oilseed rape and wheat experiments. However, the feral radish plant height was 65.4 ± 1.8 cm in the sunflower experiment. Interference of feral radish may result in indirect yield losses at harvest because it matures late in the growing season and farmers may avoid harvesting densely infested areas. The presence of mature feral radish plants at the same height as the crop inflorescences may also interfere with harvest operations and compromise the quality by contaminating the seeds with weed seeds and foreign matter, lowering the value of the harvested grains (Varanasi et al. 2016). This may be particularly important in the oilseed rape and wheat crops because the feral radish plants were taller than the crops; however, it would not be important in sunflower because the feral radish height was much lower (>50 cm) than the crop.

Feral radish dry biomass per plant and per area were very similar in oilseed rape and wheat experiments, and at least seven times higher than in the sunflower experiment (Fig. 4 A and B). Feral radish dry biomass combined over both oilseed rape and wheat experiments approximately triplicated, from about 265 – 380 g m^−2^ to 850 – 1050 g m^−2^, as weed density increased at least six-fold (from 2 to >12 plants m^−2^) (Fig. 4 B). At similar densities, feral radish dry biomass in sunflower experiment increased from about 35 to 115 g m^−2^ (Fig. 4 B). The rapid stem elongation and leaf expansion characteristic of sunflower (Presotto et al. 2017) could have shaded the feral radish plants and reduced their growth. These results were consistent with higher yield losses in oilseed rape and wheat than in sunflower. At similar weed densities, Blackshaw et al. (2002) and Eslami et al. (2006) reported slightly lower values of dry biomass in wild radish under interference with oilseed rape and wheat, respectively. Conversely, feral radish seeds per pod and seed weight were not affected by weed density in any experiment (data not shown).

**Figure 4.**
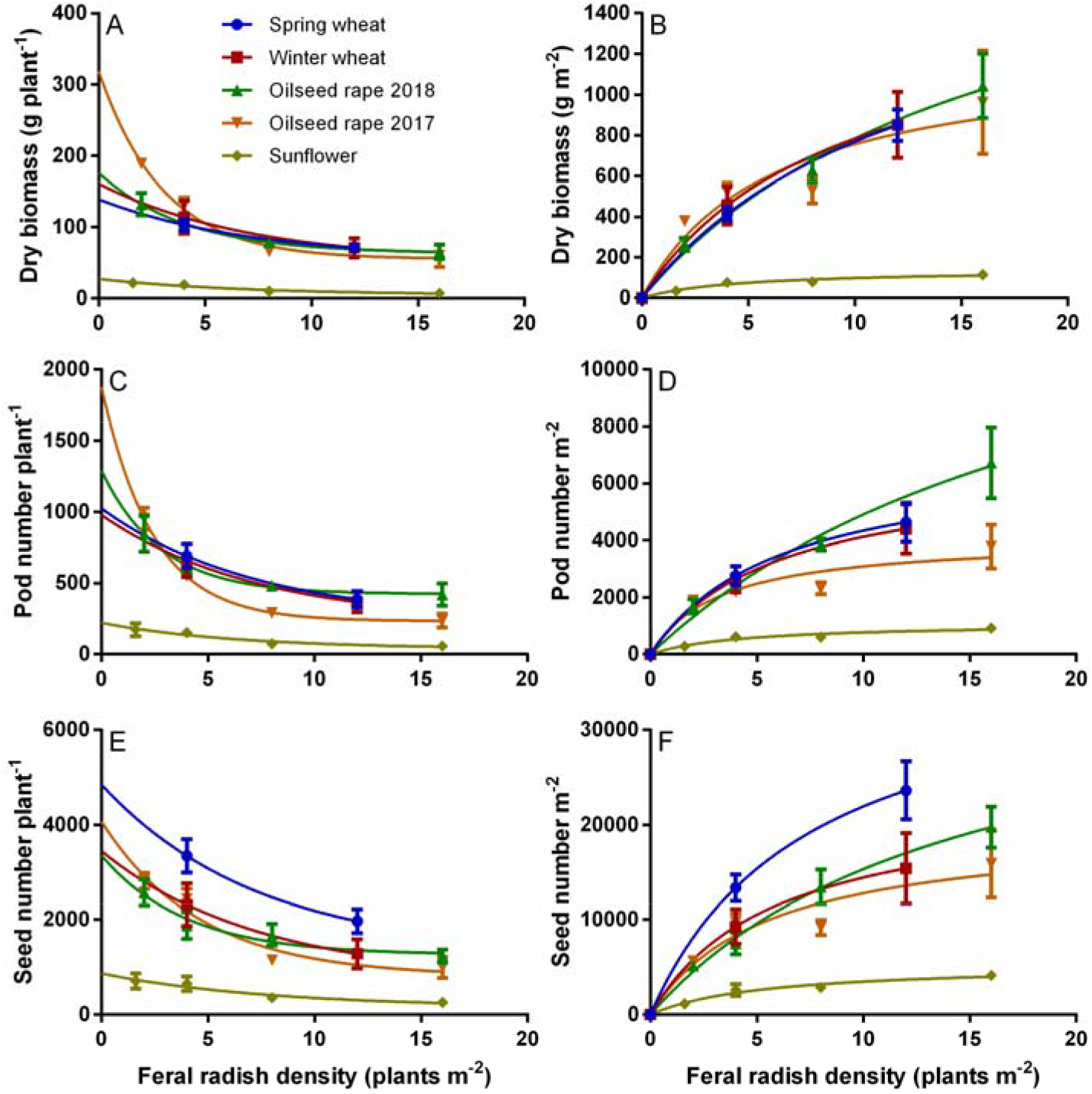
Relationship between feral radish density and dry matter per plant (A), dry matter per square meter (B), pod production per plant (C), pod production per square meter (D), seed production per plant (E) and seed production per square meter (E) of feral radish in oilseed rape (2017 and 2018), wheat (winter and spring) and sunflower experiments described with one-phase decay or hyperbola models. Vertical bars indicate means ± SE.

Preventing weed seed production in herbicide-resistant populations is a major concern among farmers around the world in order to improve weed management and/or crop production (Norsworthy et al. 2012). Feral radish pod and seed production in the oilseed rape and wheat experiments were at least four times higher than in the sunflower experiment (Fig. 4C, D, E and F). Feral radish produced around 4,300 to 6,700 and 13,800 to 31,200 seeds m^−2^ at low (2 plants m^−2^) and high (>12 plants m^−2^) densities, respectively, in both the oilseed rape and wheat experiments. Although feral radish seed production was only about 1,000 to 4,700 seeds m^−2^ at low and high weed densities in the sunflower experiment (Fig. 4F), it is a considerable amount of seeds that can enriches the seedbank. However, feral radish has high seed retention at maturity (i.e. the indehiscent pods remain firmly attached to the plant) (Snow and Campbell 2005), which could help in managing this weed through seed capture and destruction (Walsh et al. 2013).

The darkness requirement and the indehiscent pericarp may have the potential to prevent, delay and spread the germination of a cohort over an extended period (Vercellino et al. 2019), which therefore results in the formation of a persistent seedbank and hinders weed control due to the lack of synchronization in germination (Chauhan et al. 2006), what make the need to destroy the feral radish seeds/pods before to dispersion even more evident.

The importance of maintaining a weed free environment in the early growth stages of crops to prevent yield losses has been stablished in several crops (Harker et al. 2008; Page et al. 2012). Few options to control AHAS-resistant feral radish in pre- and post-emergence of sunflower, wheat and oilseed rape are effective. Since there are no chemical alternatives for pre- and post-emergence control of AHAS-resistant feral radish in oilseed rape, its sowing is not recommended in environments with presence of AHAS-resistant broadleaf weeds; at least in countries where glyphosate-resistant oilseed rape is forbidden. In sunflower, Pandolfo et al. (2016) showed effective control of AHAS-resistant feral radish with the pre-emergence mixture of acetochlor and fluorochloridone, and with aclonifen in post-emergence. In winter cereals, phenoxy (2,4-D and MCPA) and photosystem-II inhibitor (bromoxynil) herbicides are frequently used to control AHAS-resistant feral radish in post-emergence. But, the repeated use of these herbicides to control AHAS-resistant feral radish could led to the evolution of multiple-herbicide resistant populations as it has happened in wild radish (Walsh et al. 2004, Lu et al. 2019).

This study shows that the interference of feral radish can cause considerable yield losses in oilseed rape, wheat and sunflower. The high seed production, and the prolonged and irregular emergence pattern offers an adaptive potential to feral *R. sativus* to become a dominant weed. However, high seed retention at maturity provides an opportunity for controlling feral radish through harvest weed seed control tactics. The use of all cultural, mechanical and herbicidal options available for effective feral radish control must be considered in order to minimize crop yield losses and avoid/reduce weed-resistant seed production (Norsworthy et al. 2012, Bajwa 2014). Future research should be conducted to determine the interference and seed production of feral radish emerging at different times after crop sowing. These studies will provide critical information for timely management of feral radish.

## Acknowledgements

This work was supported by the grant ANPCYT-PICT 2012-2854. We thank the National Research Council of Argentina (CONICET) for a fellowship to RBV. No conflicts of interest have been declared.

