## Supplementary material for "Interference of feral radish (*Raphanus sativus*) resistant to AHAS-inhibiting herbicides in oilseed rape, wheat and sunflower crops": Tables

**Table S1.** *ANOVA* for effects of five feral radish density on oilseed rape growth and reproductive traits in two years

|  | Plant height | | Number of branches | | Pods per main inflorescence | | Seeds per pod | | Seed biomass | | Seed biomass 2017 | | Seed biomass 2018 | |
| --- | --- | --- | --- | --- | --- | --- | --- | --- | --- | --- | --- | --- | --- | --- |
| Effect | *F* | *P* | *F* | *P* | *F* | *P* | *F* | *P* | *F* | *P* | *F* | *P* | *F* | *P* |
| Year (Y) | 108.69 | <.0001 | 14.43 | 0.0009 | 18.84 | 0.0002 | 1.11 | 0.3021 | 7.84 | 0.0099 | - | - | - | - |
| Block (Y) | 2.28 | 0.0702 | 2.85 | 0.0308 | 3.25 | 0.0175 | 0.52 | 0.7853 | 3.66 | 0.0101 | 3.54 | 0.0482 | 3.78 | 0.0405 |
| Density (D) | 6.55 | 0.0010 | 18.93 | <.0001 | 6.57 | 0.0010 | 10.07 | <.0001 | 16.18 | <.0001 | 1.22 | 0.3522 | 22.61 | <.0001 |
| Y x D | 1.29 | 0.3029 | 0.20 | 0.9368 | 1.71 | 0.1802 | 0.15 | 0.9615 | 7.46 | 0.0005 | - | - | - | - |

**Table S2.** *ANOVA* for effects of three feral radish density on wheat growth and reproductive traits in two growing season

|  | Plant height | | Spikes m^-2^ | | Spikelets per spike | | Grain per spikelet | | Grain biomass | |
| --- | --- | --- | --- | --- | --- | --- | --- | --- | --- | --- |
| Effect | *F* | *P* | *F* | *P* | *F* | *P* | *F* | *P* | *F* | *P* |
| Growing season (GS) | 14.24 | 0.0027 | 54.92 | <.0001 | 0.21 | 0.6545 | 48.74 | <.0001 | 20.93 | 0.0006 |
| Block (GS) | 0.55 | 0.76 | 0.72 | 0.64 | 2.05 | 0.14 | 2.69 | 0.07 | 1.21 | 0.36 |
| Density (D) | 0.34 | 0.7190 | 49.50 | <.0001 | 11.78 | 0.0015 | 61.86 | <.0001 | 5.25 | 0.0230 |
| GS x D | 1.71 | 0.22 | 0.60 | 0.57 | 0.05 | 0.95 | 2.23 | 0.15 | 0.62 | 0.55 |

**Table S3.** *ANOVA* for effects of five feral radish density on growth and reproductive traits of sunflower growing at experimental and farm level.

| Effecs | Plant height | | Number of green leaves | | Leaf area | | Head diameter | | Seeds per head | | Seed biomass | |
| --- | --- | --- | --- | --- | --- | --- | --- | --- | --- | --- | --- | --- |
| Experimental | *F* | *P* | *F* | *P* | *F* | *P* | *F* | *P* | *F* | *P* | *F* | *P* |
| Block | 3.07 | 0.0688 | 5.44 | 0.0135 | 0.28 | 0.8373 | 28.88 | <.0001 | 7.11 | 0.0053 | 2.45 | 0.1138 |
| Density | 3.33 | 0.0473 | 4.24 | 0.0228 | 2.94 | 0.0661 | 4.96 | 0.0136 | 1.62 | 0.2321 | 0.30 | 0.8722 |

**Table S4.** Parameter estimates for functions describing the effect of feral *Raphanus sativus* (feral radish) density (D) on dry biomass, pod number and seed production per plant and per area (m^-2^) of feral radish

|  |  | Y = b + (a - b)^-cD^ | | | |  | Y = ab/(b + D) | | |
| --- | --- | --- | --- | --- | --- | --- | --- | --- | --- |
|  |  | a | b | c | r^2^ |  | a | b | r^2^ |
| Oilseed rape 2017 | Dry biomass (g plant^-1^) | 319.3 | 54.95 | 0.3322 | 0.60 | Dry biomass (g m^-2^) | 1225 | 6.19 | 0.86 |
| Oilseed rape 2018 |  | 176.2 | 62.99 | 0.2463 | 0.88 |  | 2109 | 16.86 | 0.64 |
| Winter wheat |  | 160.7 | 49.37 | 0.1365 | 0.62 |  | 1510 | 9.25 | 0.78 |
| Spring Wheat |  | 138.9 | 54.45 | 0.1365 | 0.30 |  | 1803 | 13.45 | 0.94 |
| Sunflower |  | 27.3 | 4.56 | 0.1466 | 0.59 |  | 142 | 4.46 | 0.79 |
| Oilseed rape 2017 | Pods number plant^-1^ | 1878 | 232.6 | 0.4128 | 0.58 | Pods number m^-2^ | 4093 | 3.25 | 0.82 |
| Oilseed rape 2018 |  | 1291 | 423.5 | 0.3611 | 0.91 |  | 15758 | 22.10 | 0.72 |
| Winter wheat |  | 979 | 219.4 | 0.1365 | 0.44 |  | 7016 | 6.15 | 0.86 |
| Spring Wheat |  | 1024 | 232.7 | 0.1365 | 0.61 |  | 6618 | 6.04 | 0.78 |
| Sunflower |  | 221 | 37.3 | 0.1564 | 0.50 |  | 1103 | 4.38 | 0.79 |
| Oilseed rape 2017 | Seeds production plant^-1^ | 4072 | 805 | 0.2160 | 0.55 | Seeds production m^-2^ | 19841 | 5.49 | 0.89 |
| Oilseed rape 2018 |  | 3357 | 1261 | 0.2537 | 0.81 |  | 38056 | 14.81 | 0.70 |
| Winter wheat |  | 3448 | 763 | 0.1365 | 0.63 |  | 23062 | 5.95 | 0.89 |
| Spring Wheat |  | 4852 | 1275 | 0.1365 | 0.37 |  | 38310 | 7.45 | 0.70 |
| Sunflower |  | 870 | 149 | 0.1269 | 0.44 |  | 5338 | 5.33 | 0.79 |
